# Comprehensive identification of coronary artery disease-associated variants regulating vascular smooth muscle cell gene expression

**DOI:** 10.1101/2024.10.07.617126

**Authors:** Nicolas Barbera, Lexi Wallace, Noah Perry, Lily Lei, Hester M den Ruijter, Mete Civelek

**Affiliations:** Department of Genome Sciences, University of Virginia. Charlottesville, Virginia; Department of Biomedical Engineering, University of Virginia. Charlottesville, Virginia; Laboratory of Experimental Cardiology, University Medical Center Utrecht, Utrecht University, the Netherlands

## Abstract

Coronary artery disease (CAD) is a complex disorder with genetic and environmental influences. Genome-wide association studies (GWAS) have identified over 300 genomic loci associated with disease risk. However, identifying the functional variants within these loci has been limited in large part due to linkage disequilibrium. This represents a critical step in understanding the molecular mechanisms underlying disease risk. To identify and prioritize candidate causal CAD-associated variants, we performed lentivirus-based massively parallel reporter assays (lentiMPRAs) in primary vascular smooth muscle cells (SMCs), which play significant roles in atherosclerosis, the underlying cause of CAD. We tested 25892 CAD-associated variants for their allele-specific enhancer activity in quiescent and proliferative SMCs, modeling healthy and disease conditions. We identified 122 candidate variants showing significant enhancer activity and differences in reporter gene expression between risk and non-risk alleles. We also identified 23 variants showing condition-biased allelic imbalance and 41 variants showing sex-biased allelic imbalance. We further functionally characterized 25892 variants by performing CUT&RUN assays to identify variants in enhancer and promoter regions of SMCs. By integrating the results of these experiments, we identified a credible set of 49 CAD-associated variants in functionally relevant regions. Furthermore, by comparing these identified variants with our previously obtained expression quantitative trait loci (eQTL) data, 27 of these 49 variants were associated with SMC gene expression levels. Finally, we performed CRISPRi experiments on 8 variants comprising 9 variant-gene pairs, rs35976034 (*MAP1S*), rs4888409 (*CFDP1*), rs73193808 (*MAP3K7CL*), rs67631072 (*INPP5B/FHL3*), rs1651285 (*SNHG18*), rs17293632 (*SMAD3*), rs2238792 (*ARVCF*), rs4627080 (*NRIP3*), to confirm their regulatory potential of nearby gene expression. Taken together, our results comprehensively fine-map the causal variants that confer increased risk of CAD through their effects on vascular smooth muscle cells.

## Introduction

Coronary artery disease (CAD) is the leading cause of death in the US^1^. Genetic factors strongly contribute to an increased risk of CAD, with heritability estimates varying from 40% to 70%^2^. While a strong association between genetics and CAD has been established, the mechanisms through which genetic variation contributes to CAD risk remain largely unknown. A key missing link is the connection between specific, CAD-associated single-nucleotide variants and alterations in gene expression, particularly in disease-relevant tissues. Recent genome-wide association studies (GWAS) have identified over 300 genomic loci associated with CAD^3–5^. However, identifying the causal variants within these loci remains challenging. Identification is confounded by several factors. GWAS lead variants are frequently in linkage disequilibrium (LD) with dozens or hundreds of other variants, obscuring the true causal variant in a given locus. Moreover, we previously showed that 94% of CAD-associated genetic variants are in noncoding regions, suggesting that a substantial fraction of the genetic susceptibility to CAD is due to single nucleotide variants affecting changes in gene expression^6^.

Many GWAS loci are predicted to affect the vascular wall by regulating gene expression in vascular smooth muscle cells (VSMCs)^2,7^. We and others have identified the molecular mechanisms of atherosclerosis susceptibility in several GWAS loci, which point to mechanisms that involve changes in the phenotypes of vascular smooth muscle cells^8–11^. Likewise, we previously identified 4910 expression quantitative trait loci in vascular smooth muscle cells, 85 of which colocalized with CAD loci^12^. However, despite these studies, identifying causal, CAD-associated variants regulating VSMC gene expression remains elusive.

In this study, we used a multi-omics approach that integrated orthogonal experimental methods, including lentivirus-based, massively parallel reporter assays (lentiMPRAs) and cleavage-under-targets-&-release-using-nuclease (CUT&RUN) assays in primary vascular smooth muscle cells along with assay for transposase-accessible chromatin with sequencing (ATAC-Seq) and eQTL data to comprehensively characterize and prioritize causal, CAD-associated variants and identify CAD-associated SNP-gene pairs. We performed these experiments in the quiescent and proliferative phenotypic state of SMCs as well as in male and female donors to identify gene-by-environment interactions.

## Results

### Overview of the prioritization pipeline

We hypothesized that a subset of the variants in CAD GWAS loci affects SMC function. We employed a combination of orthogonal experimental approaches to identify these CAD-associated candidate causal variants regulating gene expression in vascular SMCs. The overview of our approach is outlined in **Figure 1**. We performed experiments in aortic SMCs derived from healthy heart transplant donors, as previously described^13^. We cultured a sex-balanced set of 6 donors (3 male/3 female) in two conditions to model the quiescent and proliferative phenotype of SMCs observed in atherosclerotic lesions^14–17^. These conditions led to differential expression of canonical SMC markers (VCAM1, SMTN, ICAM1, TAGLN, CNN1, ACTA2, SPP1) in agreement with the differences observed in the quiescent and proliferative state of SMCs^12^. Our variant prioritization pipeline was as follows: in parallel, we performed both lentivirus-based massively parallel reporter assays (lentiMPRAs) and Cleavage Under Targets and Release Using Nuclease (CUT&RUN) assays in cultured SMCs^18,19^. We used the lentiMPRAs to identify candidate causal variants showing enhancer activity and significant differences in reporter gene expression between alleles (allelic imbalance). At the same time, we used CUT&RUN epigenome profiling to functionally annotate the variant list and identify those variants in active enhancer and promoter regions. These results were combined with our previously generated assay for transposase-accessible chromatin with sequencing (ATAC-seq) data to rank each candidate variant. We performed colocalization analysis with previously identified SMC expression quantitative trait loci (eQTLs) and CAD GWAS summary statistics to connect these candidate variants with their effector transcripts. From this prioritized list of SNP-gene pairs, we selected 8 SNPs to validate via CRISPR interference (CRISPRi) experiments.

**Figure 1.**
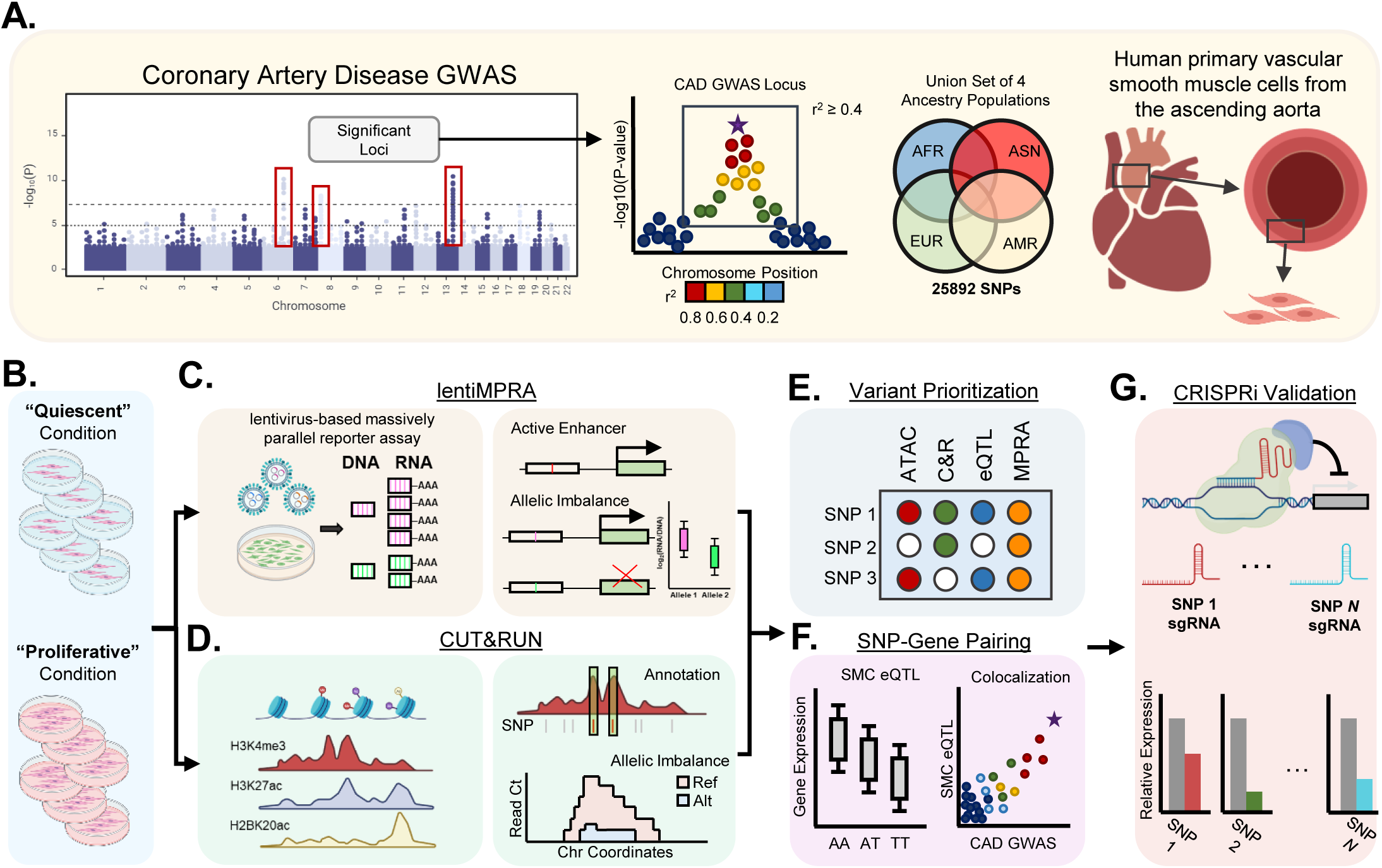
Experimental Outline. **A.** We focused on **241** loci identified in CAD GWAS ^41,42^ and identified the SNPs in moderate to high linkage disequilibrium (r^2^>0.4) in four ancestral populations. We tested their allelic enhancer activity in aortic SMCs. **B.** Primary vascular smooth muscle cells from three male and three female donors were cultured in quiescent and proliferative conditions. **C.** LentiMPRA was performed on each donor, and candidate SNPs were identified, which showed both enhancer activity and allelic imbalance. **D.** In parallel to lentiMPRA, CUT&RUN was performed on each donor for three histone modifications: H3K4me3, H3K27ac, and H2BK20ac. Peaks were identified, and candidate SNPs were annotated. Heterozygous donors were also used to identify SNPs with significantly different read counts between alleles. **E.** Candidate variants were prioritized based on orthogonal lines of experimental evidence. **F.** Prioritized SNPs were paired with genes based on CAD GWAS/eQTL colocalization data. **G.** SNP-Gene pairs were validated with CRISPRi.

### Identification of regulatory variants in CAD loci using massively parallel reporter assays

We selected the lead variants from 241 CAD GWAS loci, along with those variants in moderate to high LD (R^2^=0.4-1), based on the LD structure of four different ancestry populations (**Suppl. Fig. 1A**). In total, this yielded a set of 25982 variants, with over 97% of these existing in intronic and intergenic regions: 54.6% (14068 variants) were located within intronic regions, with another 42.4% (10918) in intergenic regions (**Suppl. Fig. 1B**). Candidate regulatory sequences (CRS) containing a 200 bp region of the genome centered on the SNP were generated for both alleles of every SNP, and cloned into a reporter assay plasmid along with unique molecular barcodes, shown in **Figure 2A**. After library packaging and sequence-barcode pairing, we found that 49134 CRS were successfully paired with unique barcodes (representing ∼95% of variants), with each CRS paired with, on average, 45 unique barcodes.

**Figure 2.**
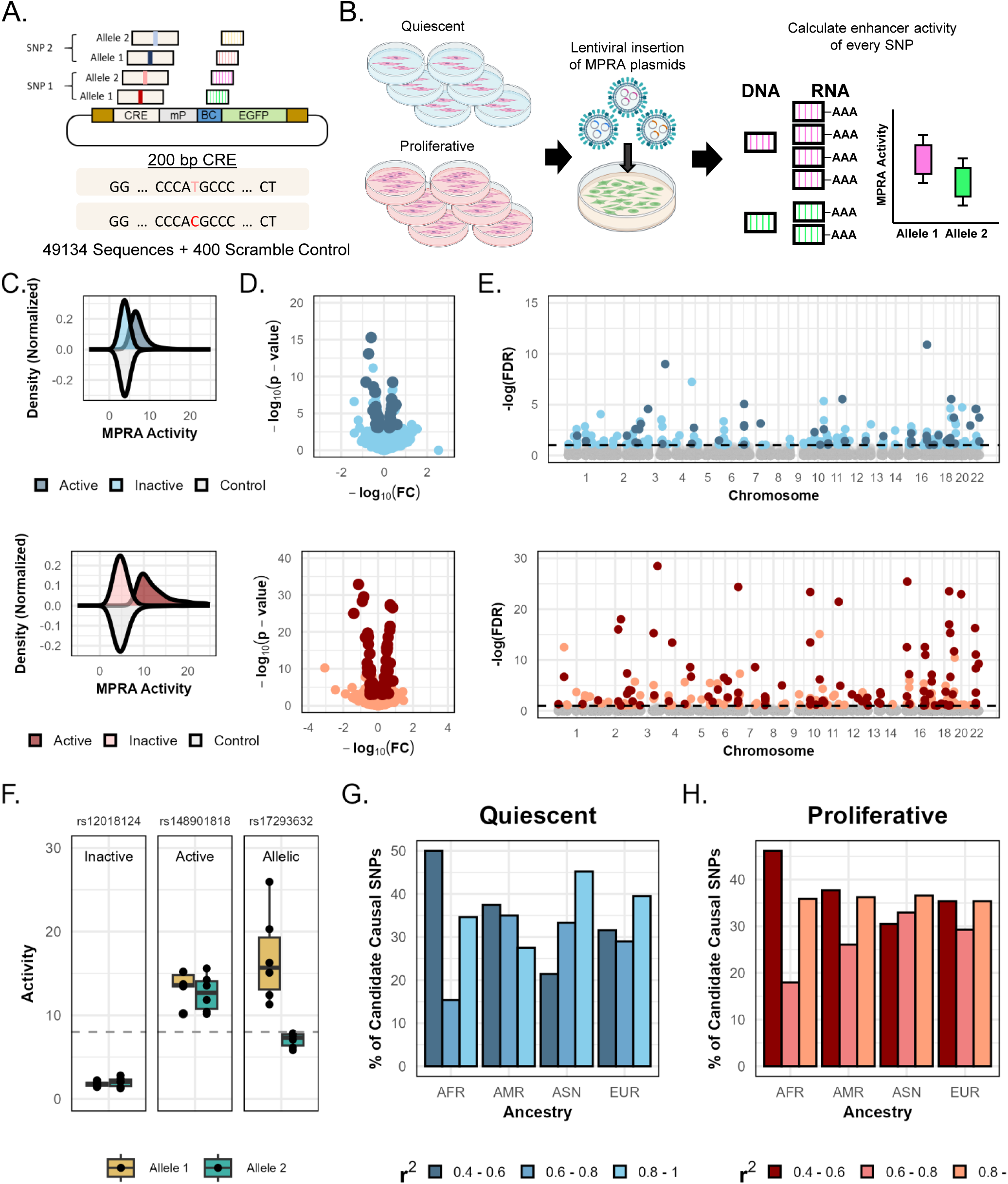
Identification of regulatory variants in CAD loci using massively parallel reporter assays. **A**. Schematic of the reporter assay plasmid used in lentiMPRA experiments. **B.** Outline of lentiMPRA experiments. Six aortic SMC donors were cultured in quiescent or proliferative phenotype conditions. A lentivirus was used to insert the MPRA plasmid into cells. After infection, RNA and DNA were simultaneously extracted and sequenced to calculate the RNA-to-DNA ratios, which were used to determine MPRA activity as a surrogate of the enhancer activity. **C.** Normalized density plots of the MPRA activity distribution for all CRS in quiescent (blue) and proliferative (red) conditions. The distribution of active sequences is shown separately in dark blue/red, while inactive and scramble control sequences are shown in light blue/red and grey, respectively. **D.** Volcano plots of the log-fold change in MPRA activity between both SNP alleles versus the calculated log normalized p-value. SNPs that contain at least one active CRS are highlighted in dark blue/red, **E.** Manhattan plots of the distribution of active and allelically imbalanced SNPs across the genome. SNPs above the dashed line showed significant allelic imbalance (FDR<0.1), while those highlighted in dark blue/red showed significant activity and significant allelic imbalance. **F.** Examples of three types of SNPs: inactive, where neither allele shows significant enhancer activity relative to scramble control; active, where there is significant enhancer activity but no allelic imbalance; and allelic, which shows significant enhancer activity and significant allelic imbalance. **G.** Bar plot showing the percentage of candidate causal SNPs identified in the quiescent SMCs in each LD bin in the four ancestry populations. **H.** Bar plot showing the percentage of candidate causal SNPs identified in the proliferative SMCs in each LD bin in the four ancestry populations.

We performed lentiMPRAs to measure the enhancer activity of CRSs simultaneously on all 6 SMC donors and in both culture conditions (**Figure 2B**). Because of the nature of library preparation, the number of barcodes associated with each CRS varied. Given that a higher number of barcodes generally leads to more accurate measures of enhancer activity, we first quantified the relationship between inter-replicate correlation and minimum barcode count. We calculated RNA/DNA count ratios for each CRS and measured the degree of correlation between donors, filtering based on the number of paired barcodes. As expected, sequences with more uniquely paired barcodes showed a greater degree of correlation in their enhancer activity across replicates, but at the cost of fewer sequences preserved for downstream analysis (**Suppl. Fig. 2**). Based on this inverse relationship, we, therefore, chose to only consider those CRS that had at least ten uniquely associated barcodes for a total of 40132 sequences out of an initial total of 51964 (77% of total CRS).

After filtering, we quantified the activity of each CRS. Using the scramble control sequences as a negative control, we identified those CRS that showed significant enhancer activity using MPRAnalyze’s quantification framework ^20^. In total, 2.6% showed activity in the quiescent condition, and 3.6% showed activity in the proliferative condition (FDR<0.1) (6.6% showed nominal significance in the quiescent phenotype and 6.8% showed nominal significance in the proliferative phenotype). The distribution of active values can be seen in the normalized density plots in **Figure 2C**. This low number of SNPs showing significant activity is comparable to what has been observed in previous MPRA experiments ^21–23^.

Having identified those sequences with significant enhancer activity, we next used MPRAnalyze’s comparative framework to identify SNPs showing significant allelic imbalance. We calculated the fold change between the MPRA activity of the two alleles of a SNP, shown in the volcano plot in **Figure 2D** along with the associated p-values. To identify candidate variants for further prioritization, we chose those SNPs showing significant allelic imbalance (FDR<0.1) and having at least one significantly active sequence between its two alleles. Following this criteria, we identified 95 SNPs in the proliferative condition and 50 SNPs in the quiescent condition for a total of 122 unique SNPs within 116 CAD GWAS loci across the genome (**Figure 2E**). Examples of active and allelic SNPs are shown in **Figure 2F**.

In addition to identifying candidate causal variants, we also set out to comprehensively characterize CAD-associated SNP activity in SMCs. We initially hypothesized that active and allelically imbalanced SNPs are over-represented in high LD with CAD GWAS lead variants. We found that this was not the case. Following the LD distribution in each 1000G population, we found that MPRA-identified candidate SNPs are distributed throughout LD thresholds and that high LD SNPs are not overrepresented (p=0.09 - 0.51) (**Figure 2G, H**).

### LentiMPRAs identify phenotype-specific SNP activity

We previously extensively characterized the culture conditions using in the lentiMPRA, including measuring significant expression differences in canonical SMC markers (VCAM1, SMTN, ICAM1, TAGLN, CNN1, ACTA2, SPP1) in agreement with the differences observed in the quiescent and proliferative state of SMCs^12,24^. Here we hypothesized that a subset of variants would show enhancer activity and allelic imbalance in a phenotype-specific manner. Therefore, we investigated the interaction between the quiescent and proliferative SMC phenotypes and variant activity. We used MPRAnalyze’s comparative framework to calculate the change in MPRA activity for each CRS between the quiescent and proliferative conditions. We found that 4240 CRS, corresponding to 3129 SNPs, showed significant differences in enhancer activity between the quiescent and proliferative conditions, suggesting that our lentiMPRA captures genotype-by-environment (GxE) effects on transcriptional activity (**Figure 3A**).

**Figure 3.**
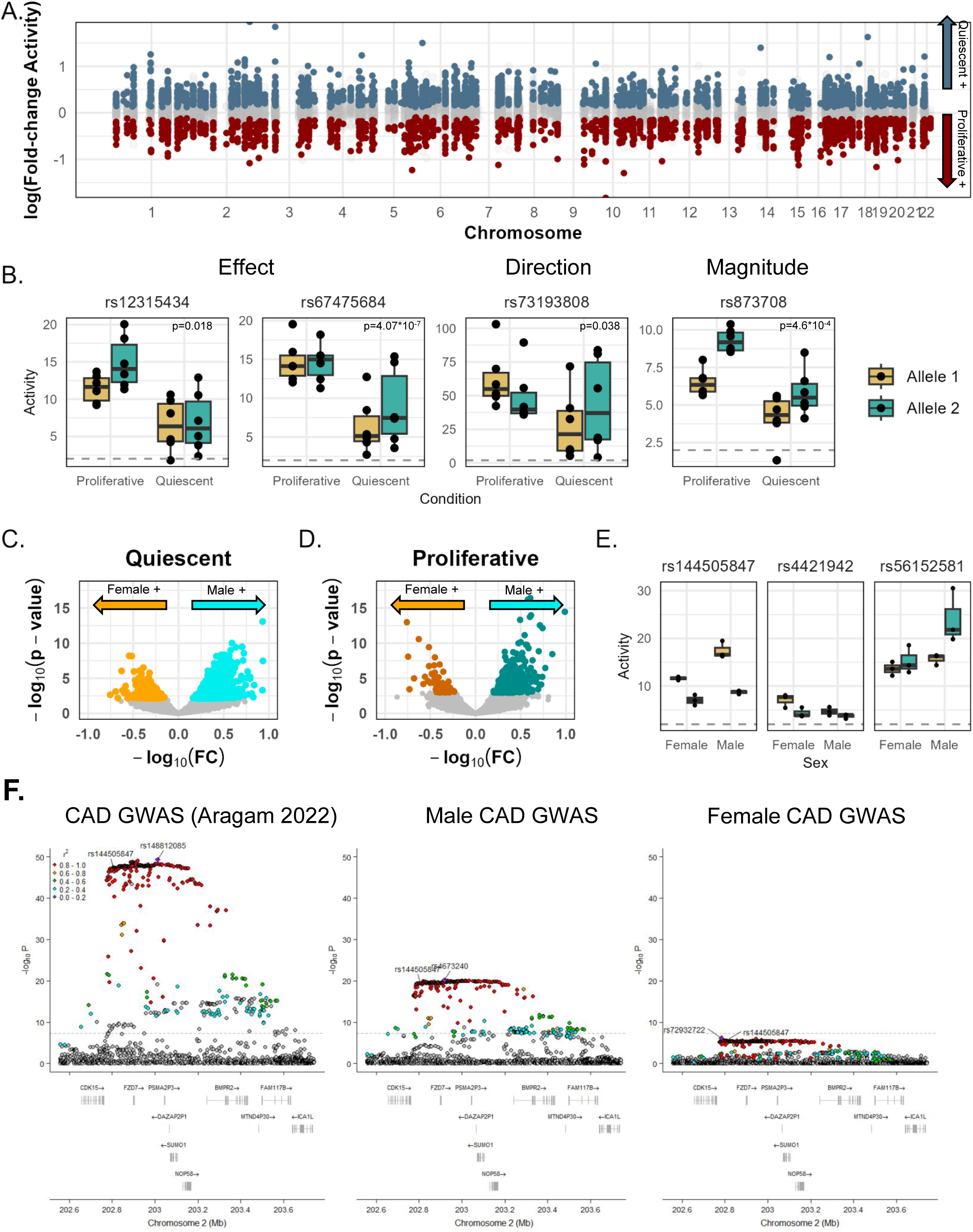
SNP-by-phenotype and SNP-by-sex effects on enhancer function. **A.** Manhattan plot of the log fold change in MPRA activity for each CRS between culture conditions. CRS highlighted in dark blue showed significantly higher MPRA activity in the quiescent condition CRS highlighted in dark red showed significantly higher MPRA activity in the proliferative condition. **B.** Examples of magnitude, effect, and directional SNP-by-phenotype effects: rs12315434 and rs67475684 show an effect difference, with rs12315434 shows an allelic imbalance in the proliferative but not quiescent condition, while rs67475684 shows an allelic imbalance in the quiescent but not proliferative condition. rs73193808 shows a directional difference in allelic imbalance, and rs873708 shows a magnitude difference. **C.** Volcano plots of the log-fold change in MPRA activity of a CRS in male vs female donors versus the calculated log normalized p-value in the quiescent condition. **D.** Volcano plots of the log-fold change in MPRA activity of a CRS in male vs female donors versus the calculated log normalized p-value in the proliferative condition. **E.** Examples of magnitude, and effect SNP-by-sex differences: rs144505847 shows a difference in magnitude of allelic imbalance. SNPs rsrs4421942 and rs56152581 show sex-dependent allelic imbalance. LocusZoom ^50^ plots showing the association signal for sex-combined (**F**) and sex-stratified GWAS for coronary artery disease at the ICA1L locus. The lead SNP rs148812085 and the identified sex-biased SNP rs144505847 are highlighted

Having found SMC phenotypic state CRS enhancer activity, we sought to identify variants showing allelic imbalance in a phenotype-dependent manner. These phenotypic differences can manifest in one of three ways: 1) a *magnitude* difference, wherein the variant shows allelic imbalance in both conditions, but the degree of allelic imbalance varies, 2) an *effect* difference, where the allelic imbalance is present in only one condition, or 3) a *directional* difference, wherein the direction of change in enhancer activity between alleles is opposite between conditions. We looked at SNPs showing significant enhancer activity and allelic imbalance in at least one phenotype to identify candidate variants. Using MPRAnalyze, we calculated an allele-by-phenotype effect. We identified 23 variants (FDR<0.1) that showed significantly different allelic imbalance between culture conditions in 23 CAD GWAS Loci. Of the 23 SNPs, 19 showed an effect difference, 4 showed a magnitude difference, and 1 showed a directional difference. Examples of SNPs showing magnitude, effect, and directional differences are shown in **Figure 3B**.

### LentiMPRAs identify sex-specific SNP activity

Sex differences exist in atherosclerosis, with females developing more fibrous atherosclerotic lesions than males ^25^. Likewise, sex differences in gene expression have been observed in many different issues and cells. In atherosclerosis, sex-based differences in plaques are mostly driven by SMCs^26^. Since we also found that there are sex-specific eQTLs in SMCs ^12^, variants may regulate expression in a sex-dependent manner. Therefore, we hypothesized sex-biased activity of candidate causal variants.

Following the same analysis approach we used to identify the candidate variants in distinct SMC phenotypes, we split our male and female grouping in both culture conditions and analyzed them separately. We found that a subset of CRS have sex-dependent activity: 530 CRS, corresponding to 427 SNPs, showed significantly different MPRA activity in 227 loci in the proliferative condition (**Figure 3C**), and 3287 CRS, corresponding to 2974 SNPs, in 240 loci showed significantly different MPRA activity in quiescent condition between male and female donors (**Suppl. Fig. 3**). We further identified 15 SNPs (FDR<0.1) that showed significantly different allelic imbalance between male and female donors in 15 CAD GWAS loci in the proliferative condition, and 41 SNPs in 33 CAD GWAS Loci showed a significantly different allelic imbalance in the quiescent condition. Examples of SNPs showing sex-dependent direction, magnitude, and effect differences are shown in **Figure 3D**.

One notable example is the SNP rs144505847 in the *ICA1L* locus. This SNP is in perfect LD (r^2^=1.0) in European ancestry populations with the index CAD GWAS variant (rs148812085). This locus was previously identified in a recent GWAS to have a between-sex heterogeneity, with higher disease risk in men as compared to women ^3^. Likewise, this SNP was shown in our MPRAs to have a greater magnitude change in enhancer activity in the male donors than female donors. Sex-stratified CAD GWAS showed distinct associations in males and females in this locus, with more significant associations in males (**Figure 3F**).

### Epigenome profiling identifies CAD variants in enhancer and promoter regions

In parallel with the lentiMPRA experiments, we performed CUT&RUN epigenome profiling assays on the same SMC donors. We focused on three histone modifications to identify CAD SNPs in enhancer (H3K27ac and H2BK20ac) and promoter (H3K4me3) regions. We used SEACR to identify genomic regions containing these histone modifications and annotated each SNP that fell within any identified peak^27^. An example of this annotation at the Chr3q.25 locus is shown in **Figure 4A**. As expected, H3K4m3e modification showed a narrow peak in the promoter region, whereas H3K27ac and H2BK20ac modifications showed a wide peak in the enhancer region of CCNL1. In this example, the GWAS signal for CCNL1 is located in the enhancer region. The lead SNP, rs4266144, is also a candidate MPRA SNP. In total, 1593 SNPs (6.1% of the CAD GWAS variants), representing 145 CAD GWAS loci, were found in at least one peak in the quiescent condition, and 3038 SNPs (11.7% of the CAD GWAS variants), representing 213 CAD GWAS loci, were found in the proliferative condition.

**Figure 4.**
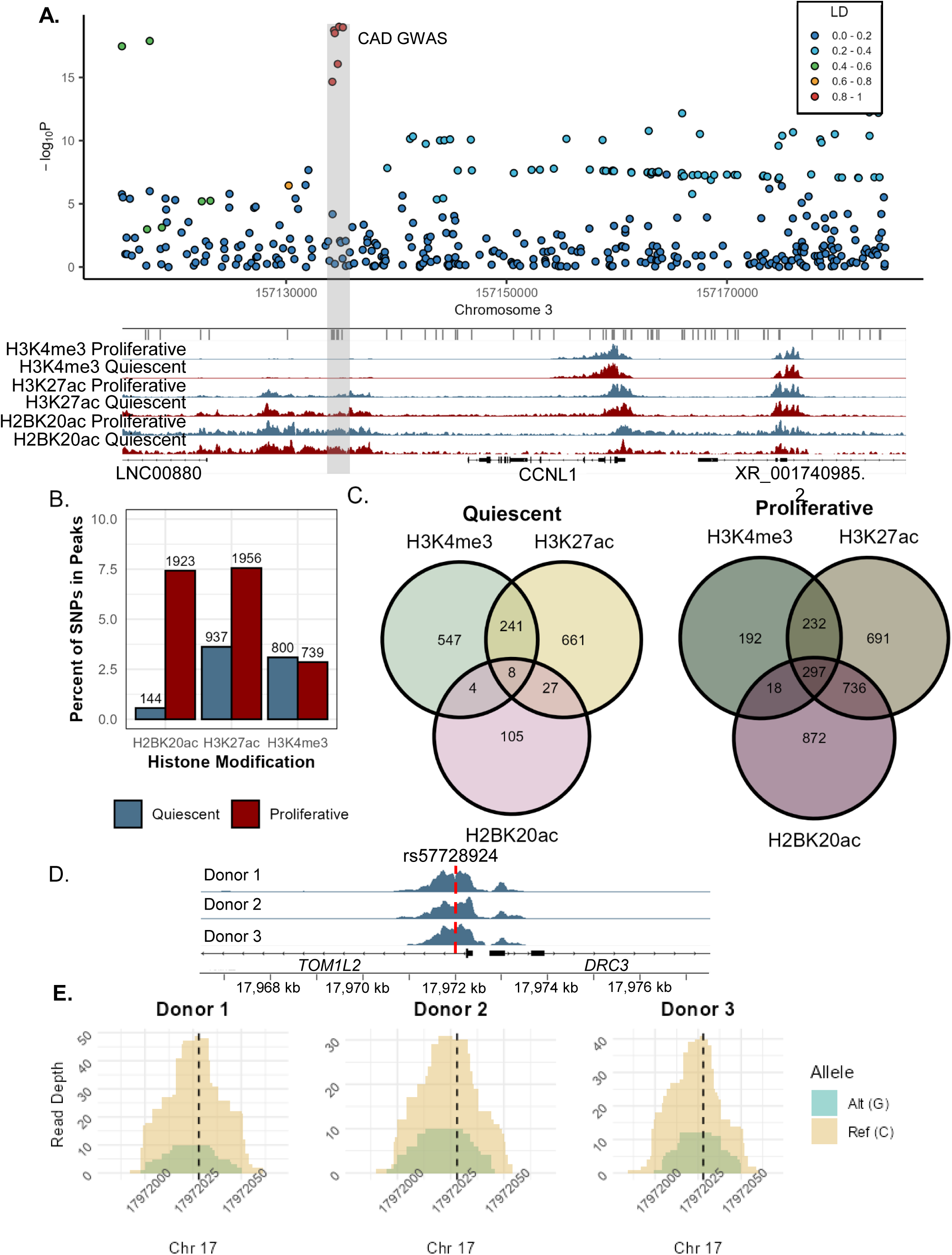
Epigenome profiling identifies CAD variants using CUT&RUN. **A.** An example of CAD loci annotation. A locuszoom plot (top) and example browser tracks of H3K4me3, H3K27ac, and H2BK20ac in quiescent (blue) and proliferative (red) conditions (bottom). The lead SNP and its associated SNPs are highlighted in the gray box. **B.** Barplot of the percentage of CAD GWAS SNPs within H3K4me3, H3K27ac, and H2BK20ac peaks in both quiescent and proliferative conditions. The number of SNPs in each type of peak is shown above the bar. **C.** Venn diagram of the overlap of SNPs found in more than one histone modification peak in the quiescent and proliferative conditions. **D.** Browser tracks of H3K4me3 in the quiescent condition in three different heterozygous donors with the location of SNP rs57728924 shown in red. **E.** Zoomed in bedgraphs of the peak shown in D, centered on rs57728924. Reads corresponding to each allele are shown separately.

Interestingly, we observed a large difference in the number of peaks between the quiescent and proliferative phenotypesfor H3K27ac and H2BK20ac but not H3K4me3, with more peaks in the proliferative cells(**Figure 4B**): SNPs (n=800) were found in H3Kme3-marked regions in quiescent cells, while 739 were identified in proliferative cells, with 671 overlapping between the two phenotypes (**Suppl. Fig. 4A**). In the quiescent cells, 937 SNPs were found in H3K27ac-marked regions, while 1956 were identified in the proliferative cells, with 733 overlapping between the two phenotypes (**Suppl. Fig. 4B**). In H2BK20ac-marked regions, only 144 SNPs were identified in the quiescent cells, while 1923 were identified in the proliferative, with 105 overlapping between the two phenotypes (**Suppl. Fig. 4C**). Furthermore, we found that within both phenotypic states, each histone modification contained a unique set of SNPs (**Figure 4C**). In the quiescent phenotype, only 280 of 1593 SNPs were shared between histone modifications, while 547, 661, and 105 were unique to H3K4me3, H3K27ac, and H2BK20ac, respectively. In the proliferative phenotype, 1283 of 3038 SNPs were shared between histone modifications, while 192, 691, and 872 were unique to H3K4me3, H3K27ac, and H2BK20ac, respectively.

In addition to functionally annotating our list of SNPs, we also used our donor set to quantify allelic imbalance in our CUT&RUN data. Of the 25982 variants we tested in our lentiMPRAs and annotated above, 2809 presented as heterozygous in at least one of the six donors used in our CUT&RUN assays. For each heterozygous SNP, we counted the number of sequencing reads that appeared for each allele and used a chi-squared test to identify SNPs showing a significant allelic imbalance between the reads mapping to the different alleles. To identify candidate causal variants, we considered only those SNPs showing significant allelic imbalance (FDR<0.1) with matching directionality in at least two separate donors in a single culture condition and histone modification. Following these criteria, we identified six candidate SNPs (**Suppl. Fig 5**): rs57728924, rs10172544, rs111465548, rs13215181,rs2258227, rs59304093. All of these appear in the H3K4me3 promoter mark, and none were identified as candidate SNPs in our lentiMPRA. The allelic imbalance in the reads overlapping the H3K4me3 promoter mark for one of these SNPs, rs57728924, was consistent across three donors (**Figure 4D & E)**.

### Integrating orthogonal datasets prioritizes 27 candidate causal variants

We hypothesized that the likeliest “true” causal variants would be those with multiple orthogonal lines of experimental support. Therefore, we combined our lentiMPRA SNPs with epigenome profiling and colocalized eQTL results. First, we considered lentiMPRA SNPs in active enhancer and promoter regions identified in our CUT&RUN data and open chromatin regions based on available ATAC-seq data from the same SMC biobank ^12,24^. We found 34 lentiMPRA SNPs in enhancer regions, 5 in promoter regions, and 27 in open chromatin regions. In total, 49 of the 122 lentiMPRA SNPs had at least one additional supporting experimental evidence (**Suppl. Fig. 6**). Additionally, we previously hypothesized that causal CAD variants act by regulating SMC gene expression. To identify the CAD GWAS loci regulating vascular wall gene expression, we performed colocalization analysis using the two most recent CAD GWAS ^3,4^ and five eQTL datasets. The eQTL datasets were aortic SMCs from our previous study ^12^, aortic artery, coronary artery, and tibial artery from the GTEx study ^28^, and coronary artery tissue eQTLs ^29^. We identified 249 genes that colocalized with a CAD GWAS locus in at least one of the eQTL datasets (**Suppl. Fig. 7**).

Of the 49 lentiMPRA SNPs 27, were in colocalized eQTLs, with 17 in aortic SMCs. To generate a final prioritized list of candidate causal SNPs, we ranked the 17 eQTL-associated based on the number of ATAC-seq and CUT&RUN annotations they possessed. A summary of the ranking results is shown in **Figure 5**. In total, we prioritized candidates in 16 CAD loci, corresponding to 21 SNP-gene pairs. An example of the overall prioritization process is shown for SNP rs35976034 and its associated gene, MAP1S (**Figure 6**).

**Figure 5.**
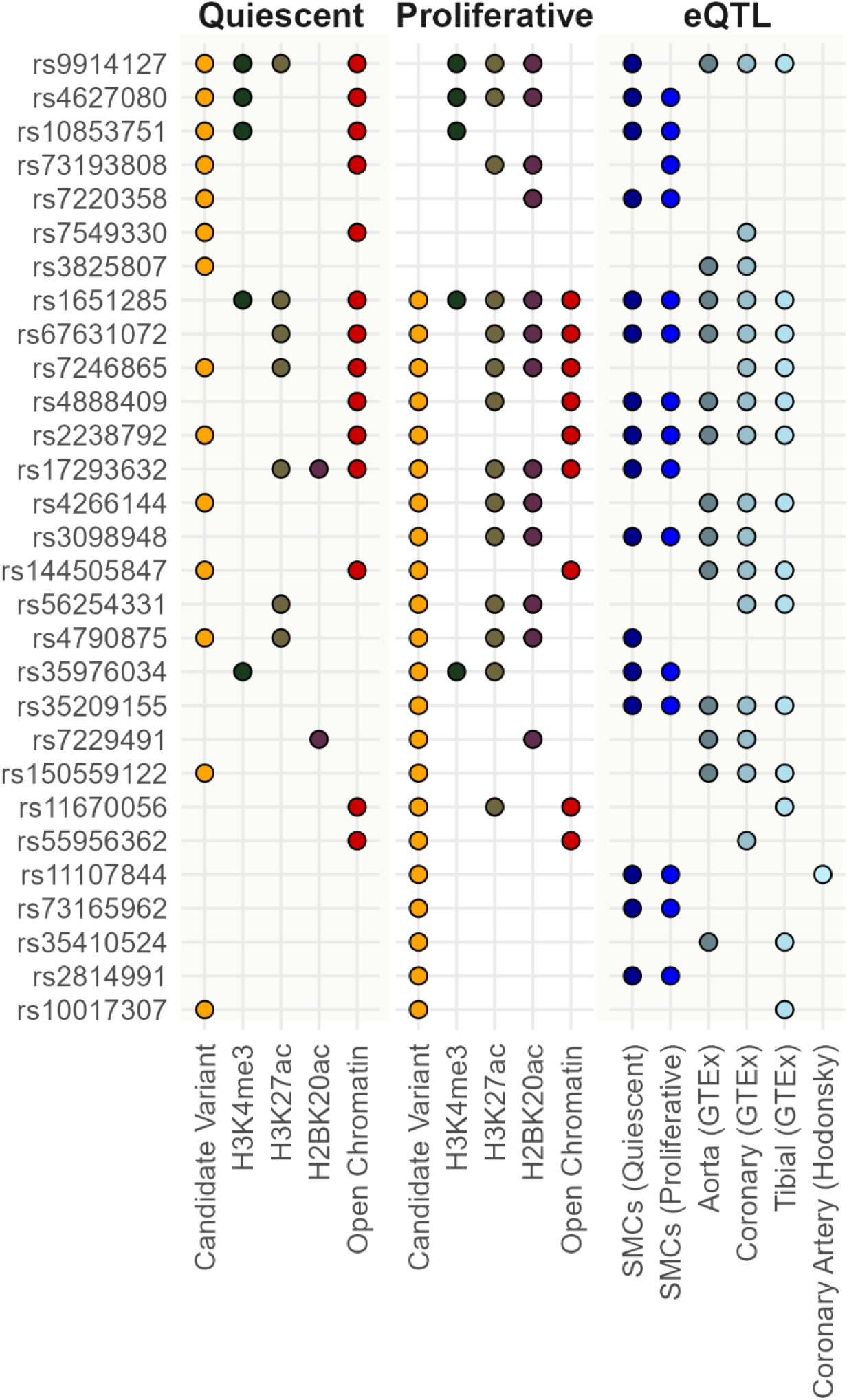
SNP Prioritization. List of the top 19 SNPs based on aggregation of experimental evidence in orthogonal assays. Colored dots denote supporting evidence for the SNP in a given assay. In the left two columns: orange denotes that the SNP was identified as a candidate SNP in the lentiMPRA, green, yellow, and purple denote that the SNP is present in H3K4me3, H3K27ac, and H2BK20ac, respectively (based on CUT&RUN), red denotes that the SNP is present in a region of open chromatin (based on ATAC-seq). In the right column, colored dots denote that the SNP is in a colocalized eQTL signal from one of eQTL datasets: SMCs in the quiescent or proliferative condition, aorta, coronary artery, or tibial artery tissue from GTEx, or coronary artery tissue.

**Figure 6.**
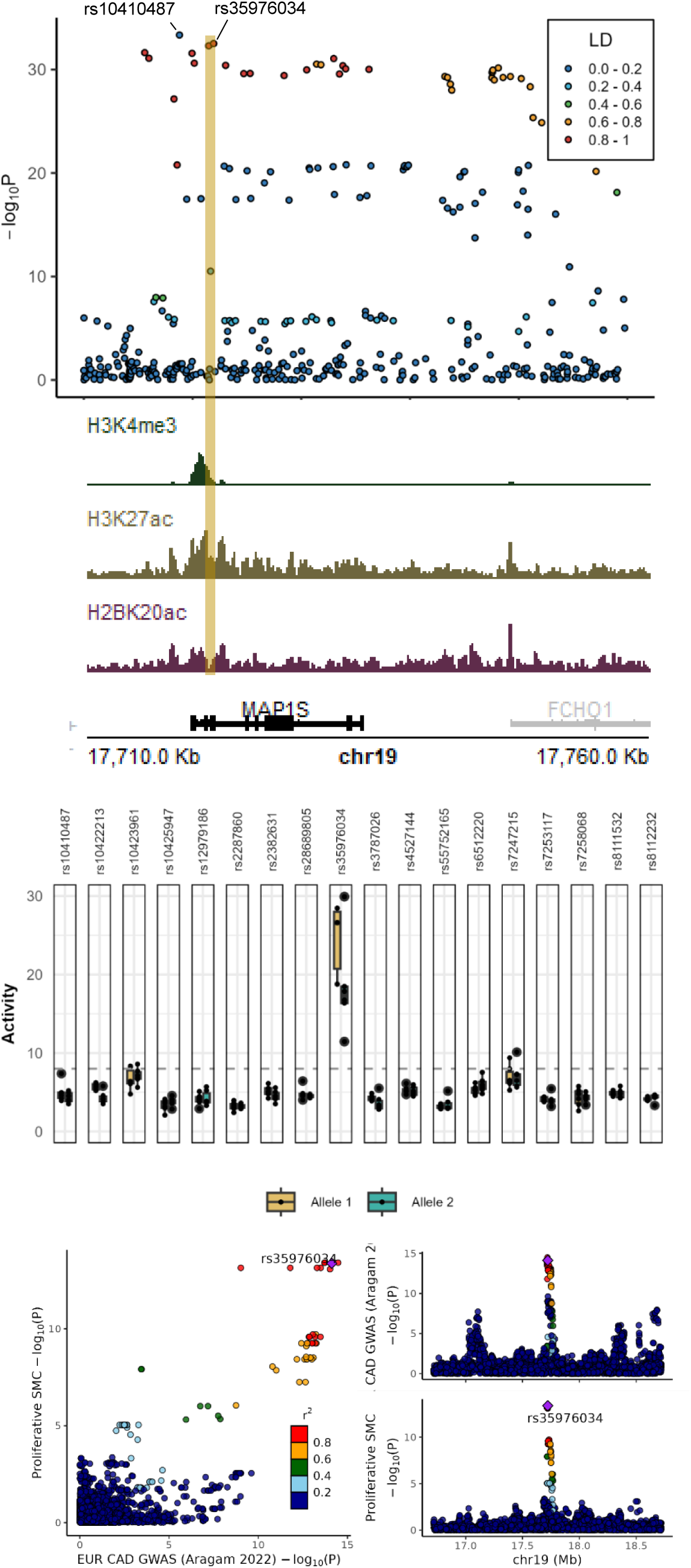
Example of Variant Prioritization. Orthogonal lines of evidence that led to the prioritization of SNP rs35976034 is shown: **A.** The CAD GWAS locus contains rs35976034. Both rs35976034 and the GWAS lead SNP (rs10410487) are labeled. **B.** Browser tracks of H3K4me3, H3K27ac, H2BK20ac for SMCs in the proliferative condition. The location of rs35976034 is highlighted in yellow. **C.** lentiMPRA results for high LD (R^2^ >0.8) SNPs tested in the GWAS locus. **D.** Colocalization results of MAP1S and the GWAS locus, with rs35976034 highlighted.

**Figure 7.**
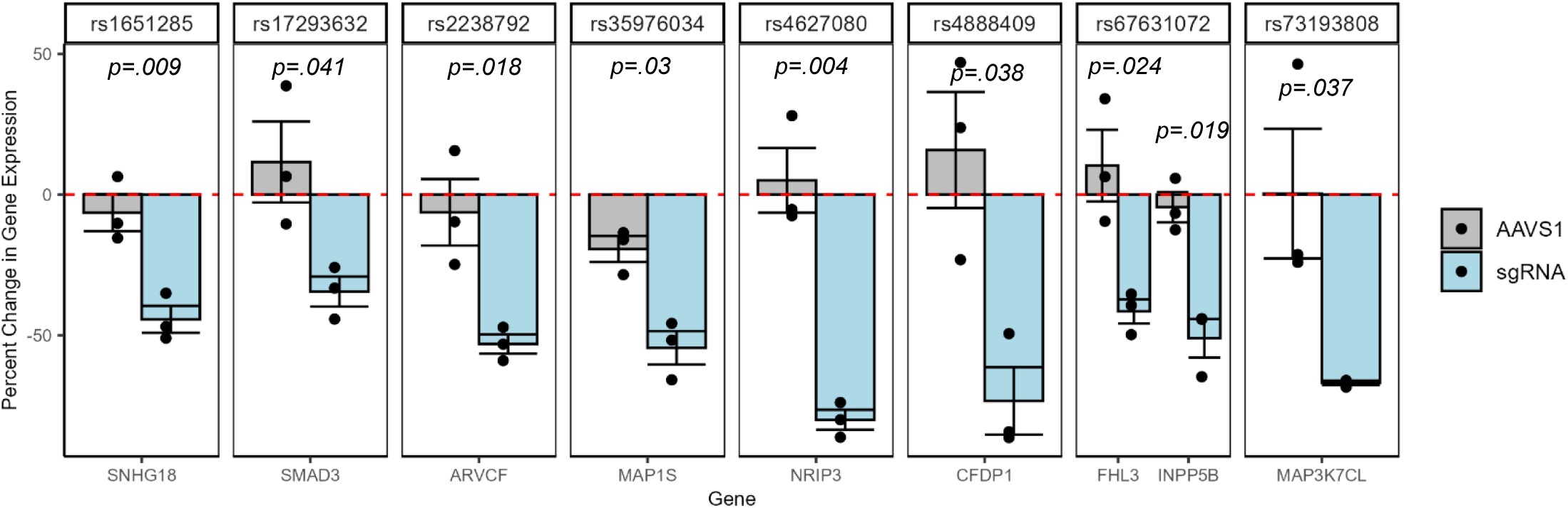
Validation of Candidate Variants with CRISPRi. CRISPRi with gRNAs targeting the regions overlapping SNPs: rs1651285 (*SNHG18*), rs17293632 (*SMAD3*), rs2238792 (*ARVCF*), rs35976034 (*MAP1S*), rs4627080 (*NRIP3*), rs4888409 (*CFDP1*), rs67631072 (*INPP5B/FHL3*), rs73193808 (*MAP3K7CL*). The levels of the respective gene transcripts (GAPDH-normalized) are shown as fold change over those from non-targeting gRNA. Error bars refer to the standard error. p values are calculated by two-sample t test (two-sided) with unequal variance from non-targeting controls (dotted red lines).

### CRISPR interference validates 8 prioritized SNPs

To validate whether the genomic regions containing our candidate SNPs regulate gene expression we performed CRISPR interference (CRISPRi) on our candidate SNP-gene pairs. For each SNP, we designed a guide RNA sequence overlapping the region containing the SNP in question. As a negative control, we also designed a guide RNA sequence to target the AAVS1 safe harbor locus. In total, we tested 8 SNPs corresponding to 9 SNP-gene pairs: rs35976034 (*MAP1S*), rs4888409 (*CFDP1*), rs73193808 (*MAP3K7CL*), rs67631072 (*INPP5B/FHL3*), rs1651285 (*SNHG18*), rs17293632 (*SMAD3*), rs2238792 (*ARVCF*), rs4627080 (*NRIP3*). All 8 SNPs are in colocalized eQTLs from our quiescent and proliferative SMC datasets, with 4 SNPs (rs1651285, rs67631072, rs4888409, and rs2238792) appearing in colocalized eQTL signals from GTEx datasets. CRISPRi followed by qPCR showed a 0.5 - 1-fold reduction in target gene levels when targeting the regions containing rs1651285, rs17293632, rs2238792, and rs67631072 vs controls. Likewise, we observed a 1.5-2-fold reduction in *MAP3K7CL, NRIP3, and CFDP1* levels when targeting the regions containing rs35976034, rs4627080, and rs488409, respectively. Overall, these data identified SNHG18, SMAD3, ARVCF, MAP3K7CL, NRIP3, CFDP1, FHL3, and INPP5B as plausible CAD susceptibility genes regulated by *cis*-regulatory variants.

## Discussion

The identification of causal variants in GWAS loci remains a challenge. Each locus contains dozens of variants in linkage disequilibrium with one another, obscuring the causal relationship between functional variants and disease risk. In this study, we deployed a multi-omics approach integrating lentiMPRAs, CUT&RUN assays, and ATAC-seq in primary vascular smooth muscle cells along with eQTL data to comprehensively characterize CAD-associated variants and identify likely causal SNPs.

The initial set of 25982 CAD-associated SNPs was generated by setting a permissive LD threshold (r^2^ >0.4) when selecting SNPs associated with CAD GWAS lead SNPs. This permissive threshold is in contrast to other recent MPRA-based studies investigating causal variants, which typically use either a higher LD threshold (r^2^ >0.8), a pre-screened set of likely variants, or both. For example, a recent MPRA study investigating blood pressure-associated variants in cardiomyocytes and stem cell-derived VSMCs used an LD threshold of r^2^ >0.8 for lead SNPs, testing around 5000 variants ^21^. Likewise, a recent study identifying variants in melanoma loci followed the same approach, testing 1992 variants ^23^, while another study investigating type 2 diabetes-associated SNPs in MIN6 beta cells tested 6621 SNPs either in high LD (r^2^ >0.8) with GWAS lead variants or residing in accessible chromatin sites in human islets ^30^. We deliberately set a permissive LD threshold to identify as many candidate variants as possible in an unbiased manner. We hypothesized that a subset of causal variants existed in moderate LD with their GWAS lead SNPs, which would be missed in a more stringent approach. Based on the results of the lentiMPRAs, this hypothesis appears correct. Depending on the ancestry population’s LD structure, between 65-68% of the 122 candidate lentiMPRA SNPs were in low-to-moderate LD (r^2^ <0.8) with their respective lead SNP. Furthermore, of the 21 causal SNP-gene pairs identified, 46-62% of SNPs were in low-to-moderate LD. These results suggest an important design consideration for future MPRAs in investigating allelic imbalance, given that many of the variants identified here would otherwise have been missed. These results also point to the necessity of using diverse ancestry populations when identifying candidate causal variants: three of our validated variants, rs67631072, rs1651285, and rs17293632, are in high linkage disequilibrium (r^2^=0.94-1) with their respective lead SNPs in the European ancestry population, while in the African ancestry population, this was not the case (r^2^=0.36-0.64).

Other recent studies have also investigated cardiovascular disease and cardiovascular trait-associated variants in smooth muscle cells. A recent study performed self-transcribing active regulatory region sequencing (STARR-seq), a type of MPRA, in human aortic SMCs and prioritized 34 CAD/myocardial infarction-associated candidate variants ^31^. However, of these 34 variants only 3, rs73193808, rs17293632, and rs1651285, were replicated in our study. One possible explanation for this difference is cell culture condition: the SMCs in this study were cholesterol loaded to induce SMC phenotype switching, whereas we did not use cholesterol in our cell culturing. Likewise another recent MPRA study investigating blood pressure-associated variants identified 84 candidate SNPs in stem cell-derived VSMCs ^21^. However, only 2 of these SNPs, rs62059846 and rs6793477, were identified as SNPs showing FDR-corrected significant allelic imbalance in our study, with another 8 showing nominally significant allelic imbalance. Both of these studies in comparison to ours highlight the importance of cell type and culture conditions in identifying regulatory SNPs.

The choice of two different culture conditions and donors from both sexes allows us to study the effects of gene-by-environment (GxE) and gene-by-sex interactions regarding causal variants. We previously identified 910 condition-specific eQTLs in SMCs ^12^. In this study, we identified SNPs showing phenotype bias in enhancer activity level and phenotype-dependent allelic imbalance, which may stem from differences in transcription factor or cofactor expression levels between the culture conditions. This is in line with what has been observed for CAD-associated SNPs in other cell types: STARR-seq experiments performed in human aortic endothelial cells showed differences in enhancer activity for SNP-containing enhancer regions when treated with IL-1β, particularly in regions containing the NF-κB motif ^32^. 5.6% of enhancer regions tested in endothelial cells treated with IL-1β had differences in activity, which is comparable to the 12% of SNP-containing regions we observed in our lentiMPRAs. With respect to the progression of atherosclerosis, this might suggest that there are different causal variants regulating gene expression - and therefore disease risk - depending on the stage of disease. In the context of SMCs, the process of dedifferentiation and phenotypic switching involves a complex interplay of transcription factors, epigenetic changes, chromatin remodeling, and changes in gene expression ^33^. Causal variants acting to regulate the expression of specific genes therefore may not be relevant until the SMCs in question have differentiated to the necessary phenotype. An example from our lentiMPRAs is the candidate SNP rs67631072. We found that the enhancer activity and allelic imbalance were only present in the proliferative condition while the gene it is associated with, FHL3, is itself associated with proliferation and transcriptional regulation ^5^. A quiescent SMC then might not be affected by this variant, and the regulatory effects of this SNP might only become a factor only after the phenotypic switching has started.

We also interrogated the question of sex differences by using a sex-balanced donor set in our lentiMPRAs. It is known that CAD risk varies between men and women ^34^. However, few studies have provided a mechanistic understanding of these differences. Sex-biased CAD GWAS loci have been identified, suggesting a genetic component to these differences and opening the possibility of using human genetics findings to provide a mechanistic understanding of the sex differences ^25^. Similar to our phenotype-dependent variants, we identified SNPs with sex-biased effects, including one SNP that is part of a known sex-biased CAD GWAS locus, rs144505847, in the *ICA1L* locus. Because these assays were done in male and female cells in identical culture conditions, these measured differences would likely come from sex chromosome-based differences rather than hormonal regulation.

While multiple studies have demonstrated their effectiveness in identifying regulatory variants, there are limitations to MPRAs. In constructing the large library used in this study, there were some restrictions. Because molecular barcodes were randomly paired with CRS in our lentiMPRA, there was a tradeoff of library complexity (barcodes paired uniquely with CRS) vs the number of testable SNPs. Due to the random process of pairing barcodes to CRSs, some SNPs excluded from this analysis based on insufficient barcode pairing could have otherwise been identified as candidate variants. Likewise, some SNPs that failed to meet the significance thresholds applied in this study would have been otherwise identified had they been paired with more barcodes in a smaller experiment. Furthermore, the large number of SNPs tested here necessitated many cells (>10 million per biological replicate) to achieve sufficient viral integration. This may not be possible for different types of primary cells.

In interpreting our results, it is also essential to consider the function of the assay itself. Due to the design of the reporter plasmid, the lentiMPRA used here tested sequences for enhancer activity. Therefore, SNPs acting as repressors or SNPs affecting promoter sequences were not considered and could represent another set of causal SNPs not identified. Moreover, while these lentiMPRAs integrate test sequences directly into the genome, the insertion is random. Therefore, the enhancer activity of the SNP in its endogenous context could not be precisely recreated in an artificial assay, and effects from regulatory mechanisms like chromatin conformation were not considered.

We compensated for the limitations of lentiMPRAs by using a multi-omics approach to prioritize identified variants. Using CUT&RUN assays, we characterized the epigenetic landscape of our SMCs and functionally annotated all of the CAD GWAS SNPs based on enhancer and promoter locations. Similar to the lentiMPRA results, we identified SNPs from over 100 GWAS loci (145-213) in enhancer (H3K27ac and H2BK20ac) and promoter (H3K4me3) regions in our SMCs. We also observed a difference in the number of SNPs identified in enhancer regions between the different culture conditions. In our proliferative condition, over a thousand additional SNPs were identified in enhancer peaks. At the same time, the number of SNPs in the H3K4me3 promoter mark was relatively unchanged, with most SNPs identified being the same between the two conditions. This suggests that the mechanisms underlying the change in the epigenetic landscape between conditions depend on changes in enhancer regions driving changes in gene expression. This is in line with a recent study, which found H3K27ac and CBP upregulation in dedifferentiating SMCs in human CAV and mouse intimal hyperplasia ^35^. Moreover, because the lentiMPRA tested SNPs for their enhancer activity, our ability to identify hundreds of CAD GWAS SNPs in enhancer regions, supports our decision to integrate multiple orthogonal datasets. These results are also in line with what has been observed in previous studies, where GWAS fine-mapped SNPs are enriched in enhancer regions ^31,36,37^

In addition to annotating CAD GWAS enhancer and promoter regions, we could also leverage the distinct genetic background of our donors to look at allelic imbalance at the SNP level. Specifically, we identified 6 SNPs showing significant allelic imbalance in their read counts for H3K4me3 in multiple heterozygous donors in both culture conditions. Importantly, these results are distinct from our lentiMPRA results. The identified allelic imbalance occurs in a promoter region for all 6 SNPs, with none showing an allelic imbalance in their enhancer activity in the MPRAs. Indeed, of the 6 SNPs, only 2 showed any enhancer activity. This suggests that the regulatory mechanism of these SNPs pertains to epigenetic regulation, rather than acting as an enhancer. For example, the SNP we highlight in the results section, rs57728924, is in the promoter region of TOM1L2, a CAD-colocalized SMC eQTL.

We hypothesized that functional variants identified affect disease risk by regulating gene expression. We therefore identified candidate SNP-gene pairs by integrating our prioritized candidate list with CAD-colocalized eQTL data in the same primary smooth muscle cells as well as CAD-relevant tissue. This approach has been used in previous studies to link candidate variants with disease-associated gene expression, including liver-specific SNPs and endothelial cells in CAD ^31,32,38^, in melanoma cell and melanocytes ^23^, and in blood pressure-associated variants in red blood cells ^21^. We identified 27 SNP-gene pairs, 17 of which were present in colocalized eQTLs from the same primary cell type as the lentiMPRA. Of note is that of the 122 candidate variants prioritized in our lentiMPRAs, only 29 had a corresponding eQTL in either SMCs or vascular tissue. Since many of the omics experiments provide sparse evidence for causal variant-to-gene-to-function pipeline, it is important to have multiple lines of evidence for a comprehensive functional evaluation of the CAD-associated variants.

We tested 8 of these SNP-gene pairs with CRISPR. The variant rs17293632 has been identified in several studies as a strong candidate variant and serves as a positive control for our prioritization pipeline ^31,39^. Likewise, rs1651285 was previously identified as a candidate variant regulating SNHG18 in our previous study ^12^. Two variants, rs4888409 and rs73193808, were previously identified as potential causal variants, however they were paired with different genes than in the present study. The variant rs4888409 was previously identified via scRNA-seq and associated with BCAR1, while our colocalization results showed a stronger association with CFDP1 ^31^. The variant rs73193808 was also previously identified in the same scRNA-seq data and found to be co-accessible with BACH1 promoters in VSMCs ^31,40^. We however observed an association with MAP3KCL. In contrast rs4627080, rs2238792, rs35976034, and rs67631072, have not been studied previously.

In conclusion, by integrating multi-omics datasets in vascular smooth muscle cells, we comprehensively characterized CAD SNPs for enhancer activity and identified a credible set of candidate causal variants.

## Methods

### Cell Culture for Quiescent and Proliferative SMCs

We previously isolated SMCs enzymatically from the explants of ascending aortas of healthy heart transplant donors at the University of California at Los Angeles (UCLA) transplant program as described previously ^13^. Here, we selected 6 donors (3 male and 3 female). We maintained the cells in Smooth Muscle Cell Basal Medium (SmBM, CC-3181, Lonza) supplemented with Smooth Muscle Medium-2 SingleQuots Kit (SmGM-2, CC-4149, Lonza) (complete media). We cultured the SMCs in complete media (containing 5% FBS) until 90% confluence. Following the same procedure as Aherrahrou et. al., to mimic the quiescent state of SMCs, we switched to serum-free media and continued to culture for an additional 24 hours. To mimic the proliferative state of SMCs, we continued to culture in complete media for an additional 24 hours ^12,24^.

### Variant selection and lentiMPRA library construction

To curate the list of variants used in our library, we first collected a list of 411 SNPs from 3 CAD GWAS studies ^41,42^, consisting of both GWAS lead SNPs and fine-mapped SNPs deemed “likely causal”. We then used HaploReg v4.1 to identify additional SNPs in moderate (R^2^ >0.4) to high (R^2^>0.8) LD with these SNPs in four different ancestry populations (AFR, AMR, ASN, EUR) from the 1000 Genomes Project. SNPs from the four populations were combined, and SNPs corresponding to indels were removed. In total, this yielded a list of 25892 SNPs.

After curating the list of SNPs, the lentiMPRA library was generated following the protocol of Gordon et al ^18^. A library of 230 bp candidate regulatory sequences (CRS) were synthesized by Agilent. Each CRS in the library corresponded to a single allele of a given SNP, corresponding to a total of 51784 sequences, along with 400 negative scramble control sequences. Each CRS was designed such that the SNP was centered at the 100th bp position in a 200 bp sequence, with an additional 15 bp of adaptor sequence on each end of the 230 bp CRS. After synthesis, two successive rounds of PCR were performed to add the minimal promoter sequence and randomized barcode sequence to each CRS in the library (**Supplemental Table 15**). The amplified PCR fragments were then cloned via Gibson assembly into the SbfI/AgeI site of the pLS-SceI vector [Addgene #137725]. The recombination products were electroporated into NEB-10beta electrocompetent cells and plated on ampicillin plates. Library assembly was verified by Sanger sequencing 16 randomly chosen colonies from the resulting plates. The plasmid library was extracted from these plates using a Qiagen Plasmid Plus Midi Kit. The number of ampicillin plates used in extraction was chosen to correspond to an average of 50 barcodes per sequence.

To associate the randomized barcodes (BCs) to their corresponding CRS, we performed PCR on a sample of the extracted library to add Illumina P5/P7 flowcell sequences (**Supplemental Table 15**) and sample indices. After PCR, the prepared sample was sequenced using custom primers (Read 1, pLSmP-5bc-seq-R1; Index1 (UMI read), pLSmP-UMI-seq; Index2, pLSmP-5bc-seq-R2; Read 2, pLSmP-bc-seq, **Supplemental Table 15**) on an Illumina NextSeq (paired end sequencing 150bp x 150bp) with a read depth of ∼120 million. After sequencing, we generated Fastq files with bcl2fastq v2.20 (--no-lane-splitting --create-fastq-for-index-reads --use-bases-mask Y*,I*,I*,Y*). CRS-BC pairing was performed using MPRAflow’s association function ^20^ with the command nextflow run association.nf --fastq-insert [R1.fastq.gz] --fastq-insertPE [R2.fastq.gz] --fastq-bc [I1.fastq.gz] --fastq-bcPE [I2.fastq.gz] --design “design_library.fa” *--cigar 200M* and *--mapq 0*.

### lentiMPRA viral transduction and sequencing

The CRS plasmid library was packaged into lentivirus using pMD2.G and psPAX2 (addgene #12259 and #12260) in HEK293T cells following the same published procedure as in library construction ^18^. Because of the complexity of the library (>5 million unique barcode-sequence pairs), 6 T225 flasks were seeded with 12 million HEK293T cells each and incubated for two days with the plasmids and titerboost reagent. After incubation, viral supernatant was filtered through polyethersulfone 0.45 µm and concentrated with Lenti-X concentrator [Takara Cat#: 631231]. The supernatant mixture was incubated for 24 hours at 4°C, then centrifuged for 45 min at 1500g and 4°C. The viral pellet was resuspended in DPBS. Viral titer was determined via qpcr^18^.

Six SMC donors (3 male/3 female) at passages 5-6 were cultured in the quiescent or proliferative conditions described above to an average count of 10 million cells per donor and condition. Lentivirus containing the MPRA library was added, corresponding to an MOI of 100, and incubated for 48 hours. After 48 hours, DNA and RNA were simultaneously extracted using a Qiagen AllPrep DNA/RNA Mini Kit. PCR library preparation was performed to add unique molecular identifiers (UMI) and Sample Index barcode sequences. We then performed 25×25bp paired-end sequencing using Illumina NextSeq 2000. We sequenced our DNA sample library to a depth of 10x and our RNA sample library to a depth of 30X for a total of 5 billion reads. Because of the short length of the amplicons (16 bp), we used a 40% phiX spike-in to preserve library complexity.

For each of the indexed DNA and RNA libraries, we demultiplexed the sequencing run and generated Fastq files with bcl2fastq v2.20 as before (parameters “--barcode-mismatches 2 --sample-sheet SampleSheet.csv --use-bases-mask Y*,Y*,I*,Y* --no-lane-splitting --minimum-trimmed-read-length 0 --mask-short-adapter-reads 0”). These commands split the sequencing data into paired-end read files and UMI file for each DNA or RNA replicate sample. To obtain DNA and RNA counts, we used the count utility of MPRAflow 1.020 (run as: nextflow run count.nf --e “experiment.csv” –design “designed_library.fa” --association “filtered_coords_to_barcodes.pickle”), where “filtered_coords_to_barcodes.pickle” is the CRS-BC pairing file generated during association sequencing.

To determine which CRS showed enhancer activity, we used MPRAnalyze’s ^20^ quantification function and calculated the transcription rate alpha for each CRS (rnaDesign = ∼ 1 and dnaDesign = ∼ replicate+barcode). Significant enhancer activity, characterized by a high transcription rate, was calculated using the 400 scrambled oligos as a baseline against which to measure the expression level of a given CRS. The quantification function calculated a mean absolute deviation (MAD) score-based p-values for each CRS, which we corrected using the Benjamini-Hochberg method to generate a MAD score-based expression false discovery rate (FDR). For each SNP in both conditions, we looked at both alleles and assigned a CRS as “active” if it had an FDR ≤ 0.05 in at least one sequence.

To measure the allelic imbalance of each active SNP, we used MPRAnalyze’s comparative function. MPRAnalyze’s comparative function uses a nested linear model that incorporates information across all the barcodes for both alleles of a given sequence, along with information from all replicates. For the allelic comparison model, we used the following designs: rnaDesign = ∼ replicate + allele, dnaDesign = ∼ replicate + barcode + allele, reducedDesign = ∼ 1. We extracted the log fold change in expression between alleles for each CRS, along with the corresponding p-value. We corrected for multiple testing with the Benjamini-Hochberg method to generate an FDR for each sequence. We defined a SNP as having an allelic imbalance if it met the cutoff of FDR ≤ 0.05. We performed this analysis separately for each culture condition. Additionally we performed this analysis separately for both sexes in both culture conditions.

To measure the effects of both condition and donor sex on allelic imbalance, we used the same comparative function from MPRAnalyze as above, but with an interaction term added to the design. For the “condition by allele” analysis, we used rnaDesign: dnaDesign:, reduced design. For the “sex by allele” analysis, we used rnaDesign: dnaDesign:, reduced design:.

### CUT&RUN Assay and Sequencing

Cleavage Under Targets and Release Using Nuclease (CUT&RUN) sequencing was performed using the assay kit from Cell Signaling Technology (cat# 86652). Briefly, SMCs were cultured in the same manner as the lentiMPRA assay in both quiescent and proliferative conditions, to an average of 1 million cells per plate. Cells were harvested with trypsin and washed with a wash buffer (20 mM HEPES-NaOH pH 7.5, 150 mM NaCl, 0.5 mM spermidine, and protease inhibitor cocktail). Cells were permeabilized with digitonin and incubated with activated concanavalin A magnetic beads. Cell-bead conjugates were then resuspended in 100 µL of digitonin buffer (wash buffer with 2.5% digitonin solution) containing 2 uL of Anti-Tri-Methyl-Histone H3 (Lys4) (CST, #9733), 1 uL of Anti-Acetyl-Histone H3 (Lsy 27) (CST, #8173), 2 uL of Anti-H2BK20ac (#34156), or rabbit IgG (CST, #66362), rotated overnight at 4°C, resuspended in 250 µL of antibody buffer and 7.5 µL of the pAG-MNase enzyme (#57813, CST), followed by the rotation at 4°C for 1 h. After MNase-induced cleavage, genomic DNA fragments were extracted using DNA spin columns (CST, #14209). Samples were prepared for sequencing with the DNA Library Prep Kit for Illumina (CST, #56795) combined with Multiplex Oligos for Illumina® (Dual Index Primers) (CST, #47538). The adaptor was diluted 1:125 to avoid contamination. The PCR enrichment step ran 20 cycles to amplify the adaptor-ligated CUT&RUN DNA.

### CUT&RUN Data Analysis

Trim Galore was utilized to remove contaminant adapters and for read-quality trimming (https://www.bioinformatics.babraham.ac.uk/projects/trim_galore/). Paired-end reads were aligned to the human reference genome (HG38) using Bowtie2 (v2.2.6) (--local --very-sensitive --no-unal --no-mixed --no-discordant --phred33 -I 10 -X 700)^43^. We likewise calculated spike-in mapping rates in each sample, aligning to yeast genome R64-1-1 and calculated the corresponding scale factor (--local --very-sensitive-local --no-unal --no-mixed --no-discordant --phred33 -I 10 -X 700 --no-overlap --no-dovetail). Picard MarkDuplicates tool was utilized to mark and remove PCR duplicates in each sample, and reads with less than 20 MAPQ score were filtered. Filtered bam files were converted to bed files with bedtools bamtobed and regions longer than 1000 bp were removed. ENCODE blacklisted regions on HG38 were then removed (https://sites.google.com/site/anshulkundaje/projects/blacklists). SEACR was used to identify significant peaks with the parameters: “norm” and “stringent.” To visualize the signal in each sample, bedtools genomecov was used to convert bed files into bedgraph files using the scale factor from the spike in step above. Integrative Genomics Viewer (IGV) was used to visualize the bigwig files^44^. We used bedtools annotate to identify CAD-associated SNPs within peaks identified by SEACR. In addition to peak identification, we examined whether CAD-associated SNPs showed allelic imbalance in their bam reads. For each SMC donor, we identified those SNPs for which the donor was heterozygous. We used the R package AllelicImbalance^45^, to count bam reads corresponding to each allele for a given SNP. A Chi-squared test was performed to determine whether the difference in read counts was significant. Multiple testing correction was performed using the Benjamini-Hochberg method. We treated SNPs that showed significant allelic imbalance and matching directionality in at least two donors as candidate causal variants.

### COLOC analysis

To determine colocalization of eQTL and CAD GWAS data, we implemented Bayesian Colocalization Analysis using Bayes Factors (COLOC) using the R package COLOC ^46^. We first selected SNPs in each CAD GWAS locus with genome-wide significance, and then created 200 KB windows around significant SNPs. Following this, we merged nearby windows (>100 KB distance) together to form loci. We input these loci into COLOC with the default priors (p1/p2 = 1 × 10^-4^, and p12 = 1 × 10^-5^), and considered a locus colocalized if PPH4, the hypothesis of a single shared causal variant for both traits within a window, was greater than 0.50. We then plotted and visually inspected all analyzed loci using LocusCompare ^47^. Loci that passed both visual inspection and colocalization criteria were considered colocalized. In total, 5 eQTL data sets were used: 3 were from GTEx (Coronary artery, aorta, and tibial artery), 2 eQTL data sets were from our previous study in SMCs, and 1 data set was from expression data in coronary artery tissue ^12,28,29^. For all 5 eQTL datasets, we performed colocalization using the most recent CAD GWAS studies ^4,48^. We also included colocalized eQTLs identified in our previous study ^12^.

### CRISPR Interference Experiments

CRISPR interference (CRISPRi) was performed in the primary vascular smooth muscle cells. Guide RNAs for 8 variants (rs1651285, rs17293632, rs2238792, rs35976034, rs4888409, rs73193808, rs67631072, rs4627080) were designed to target the genomic region overlapping the variant. Guide RNA target sites were identified using the sgRNA Scorer 2.0 algorithm ^49^. Sequences for gRNAs are listed in Suppl. Table 13. A plasmid vector for each sgRNA was cloned by VectorBuilder, and contained a GFP selection marker. Plasmids encoding gRNA or dCas9-KRAB-ZIM3 (Addgene 154472) were co-transfected into HEK293T cells along with EndoFectin (Genecopoeia LT002-03) and psPAX2 and pMD2.G packaging vectors at a ratio of 3.3:2.2:1.1 ug sgRNA/dCas9:psPAX2:pMD2.G. After incubation for three days, supernatant was collected, filtered with 0.45um filter, and Lenti-X concentrator was added at a 3:1 supernatant to reagent ratio. Viral supernatant mixture was stored at 4 °C for 24 hours, then spun down for 45 min at 1500g and 4 °C. Viral pellets were re-suspended in PBS. SMCs were cultured to ∼70% confluency before being infected with lentivirus and incubated for 48 hours. We used 3 replicates per sample. Total RNA was isolated using RNeasy kit (Qiagen) and cDNA was generated via reverse transcription using SuperScript III Reverse Transcriptase (Thermofisher) and an oligo dT primer. Gene expression levels were measured via qPCR, with 3 technical replicates per sample, using the delta-delta-CT method, normalized to GAPDH, to determine expression level.

