## Supplemental Figures for "Comprehensive identification of coronary artery disease-associated variants regulating vascular smooth muscle cell gene expression"

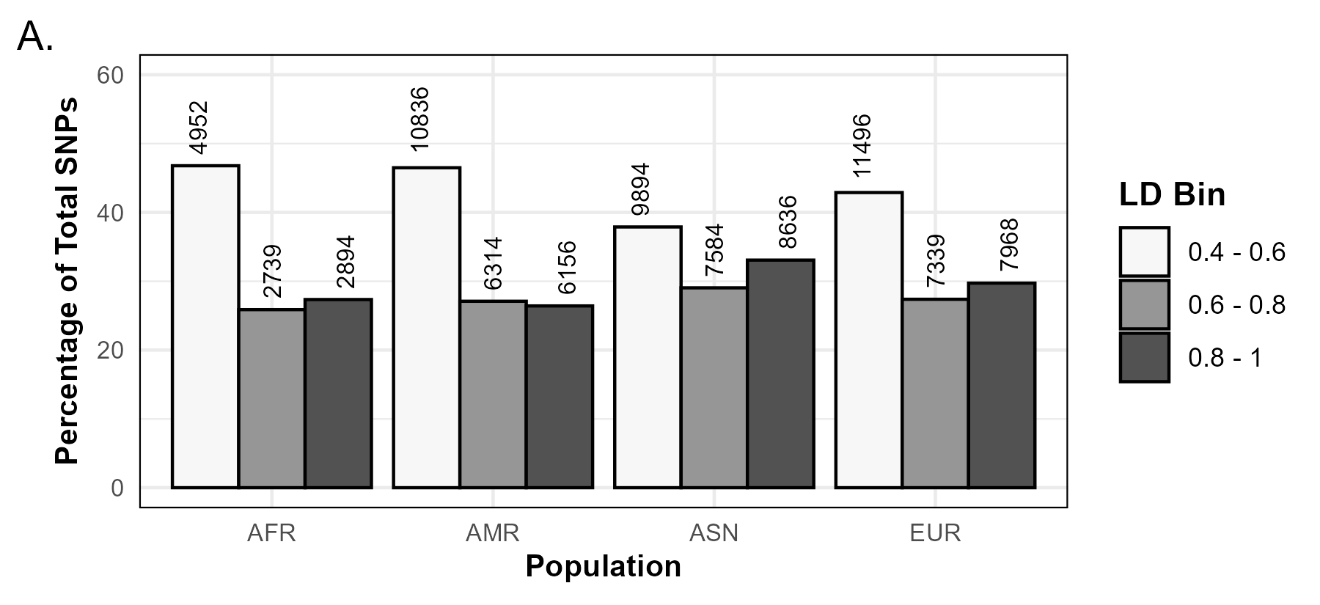

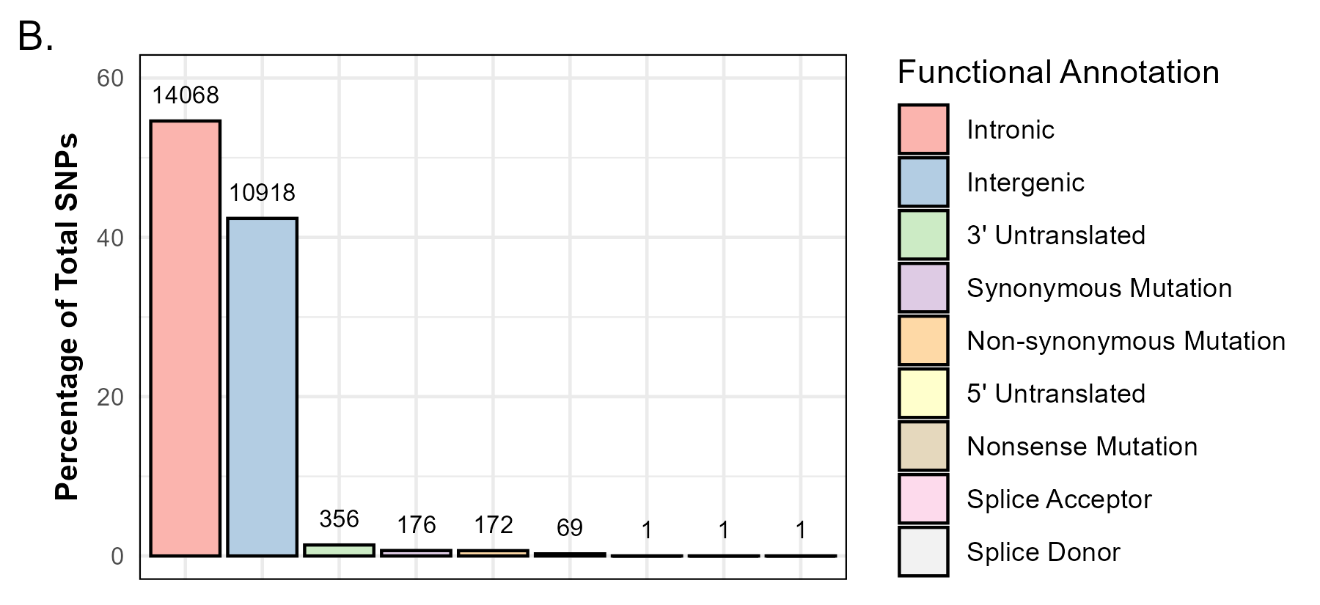
**Supplemental Figure 1. Annotating tested SNPs. A.** Bar plot showing the percentage of SNPs in each LD bin in the four ancestry populations. The number of SNPs in each LD bin is shown above each bar. **B.** Bar plot showing the percentage of SNPs with each functional annotation. The number of SNPs is shown above each bar.

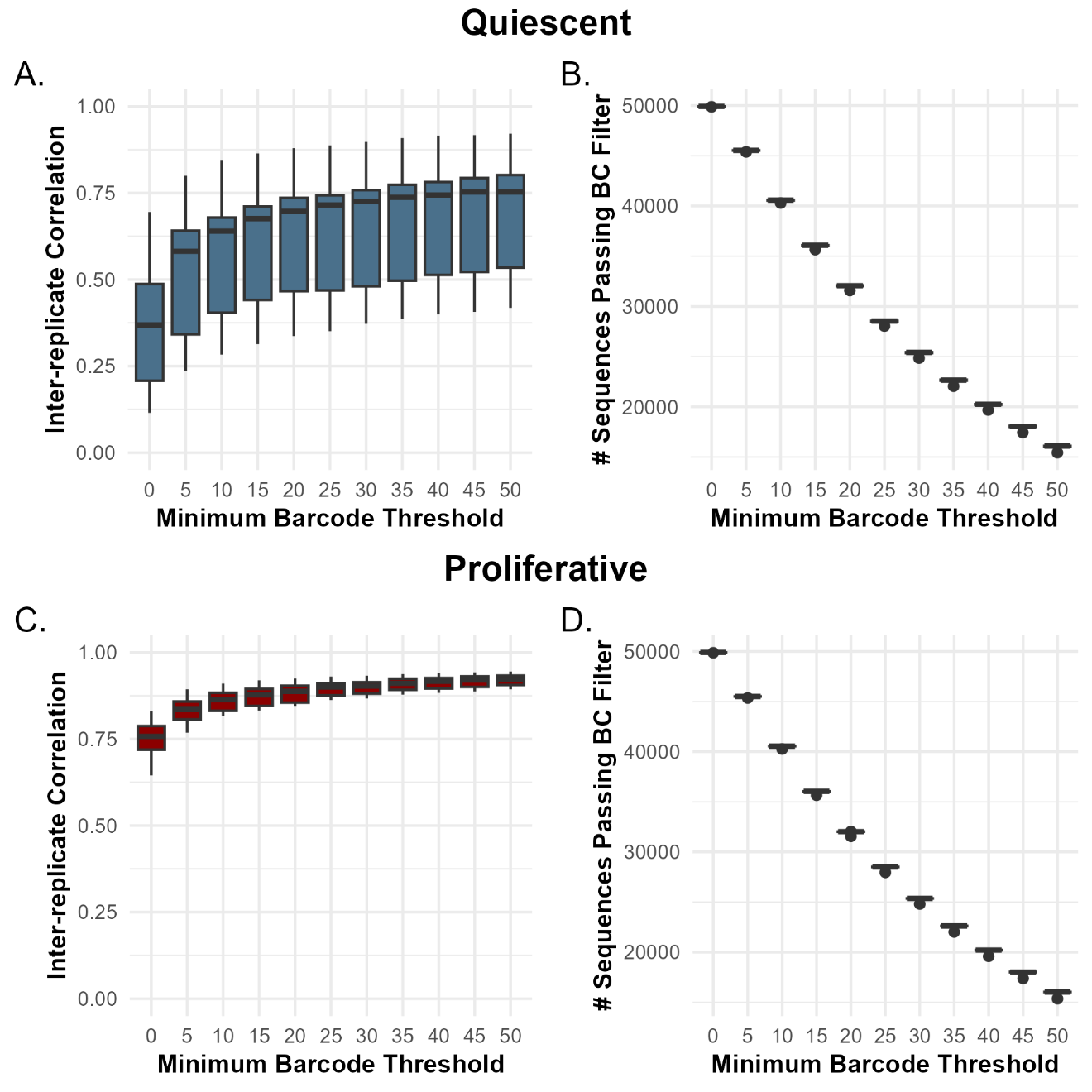

**Supplemental Figure 2.** Relationship between the inter-replicate correlation of DNA/RNA count ratios and minimum barcode threshold in the quiescent (**A**) and proliferative (**C**) conditions. Each boxplot represents the distribution of inter-replicate correlations when the results are filtered based on a minimum unique barcode threshold. Boxplots of the number of sequences in each replicate after applying barcode filtering in the **B.** quiescent and **D.** proliferative conditions.

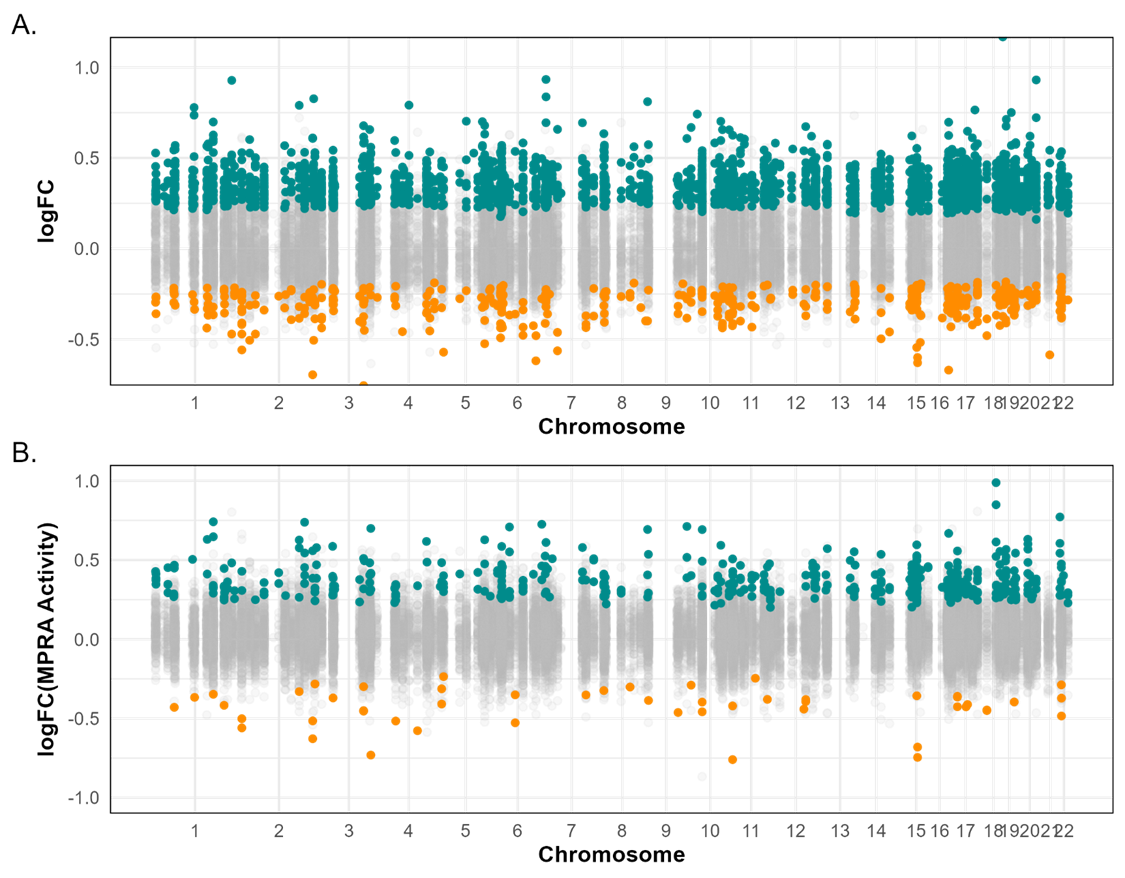

**Supplemental Figure 3. Distribution of CRS showing sex-biased activity across the genome.** Manhattan plot of the log fold change in MPRA activity for each CRS between male and female donors in the **A.** quiescent and **B.** proliferative conditions. CRS highlighted in cyan are significantly more active in the female donors, while CRS highlighted in orange are significantly more active in the female donors.

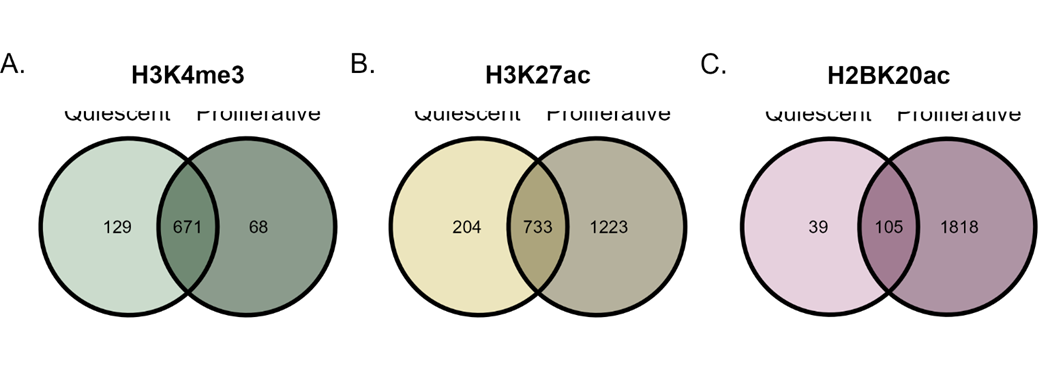

Quiescent	Proliferative

Quiescent	Proliferative

Quiescent	Proliferative

**Supplemental Figure 4. Overlap of SNPs in histone modifications by culture condition.** **A.** Venn diagram of the overlap of SNPs found in both quiescent and proliferative conditions for H3K4me3. **B.** Venn diagram of the overlap of SNPs found in both quiescent and proliferative conditions for H3K27ac. **A.** Venn diagram of the overlap of SNPs found in both quiescent and proliferative conditions for H2BK20ac.

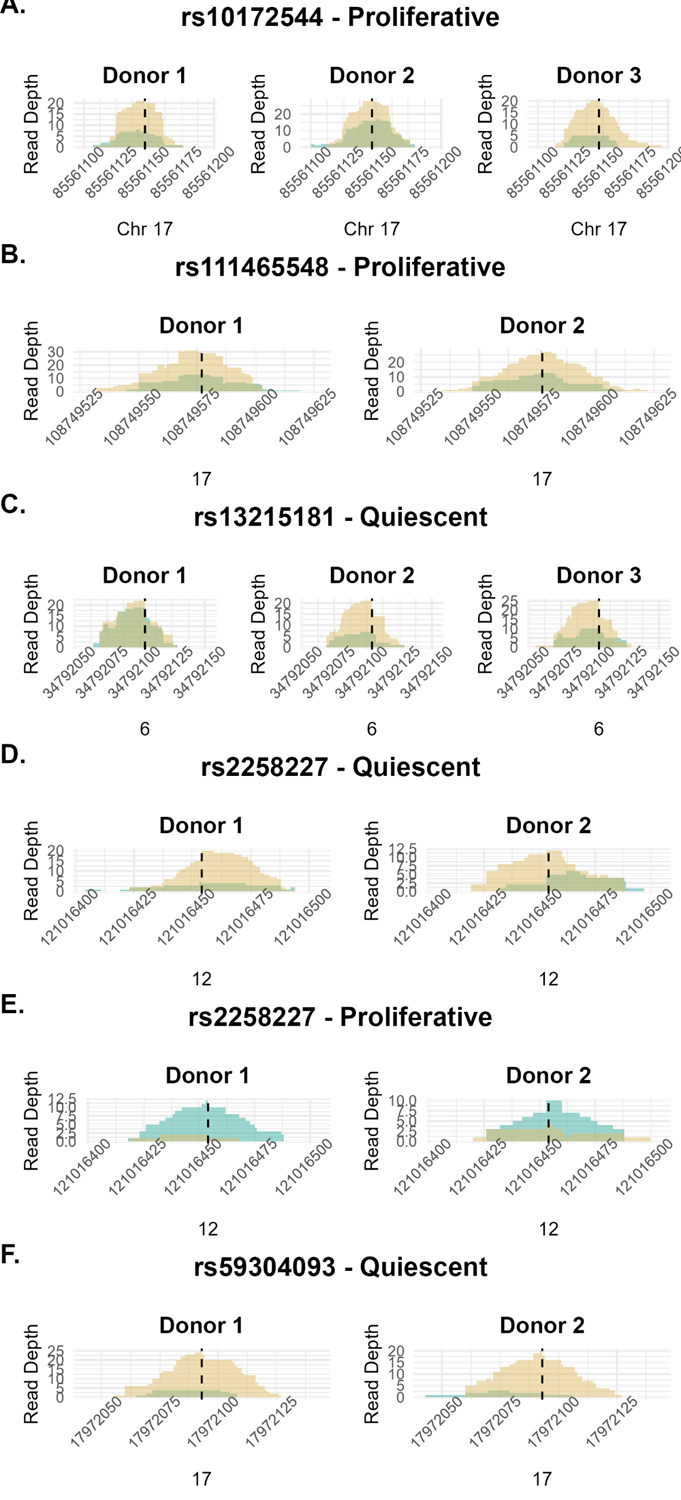

**A.**

**Supplemental Figure 5. CAD SNPs showing allelic imbalance in CUT&RUN**. Zoomed in bedgraphs of peaks for SNPs **A.** rs10172544 , **B.** rs111465548, **C.** rs13215181,  **D. & E.** rs22582227, **F.** rs59304093. Reads corresponding to each allele are shown separately. Reads from each donor are plotted separately.

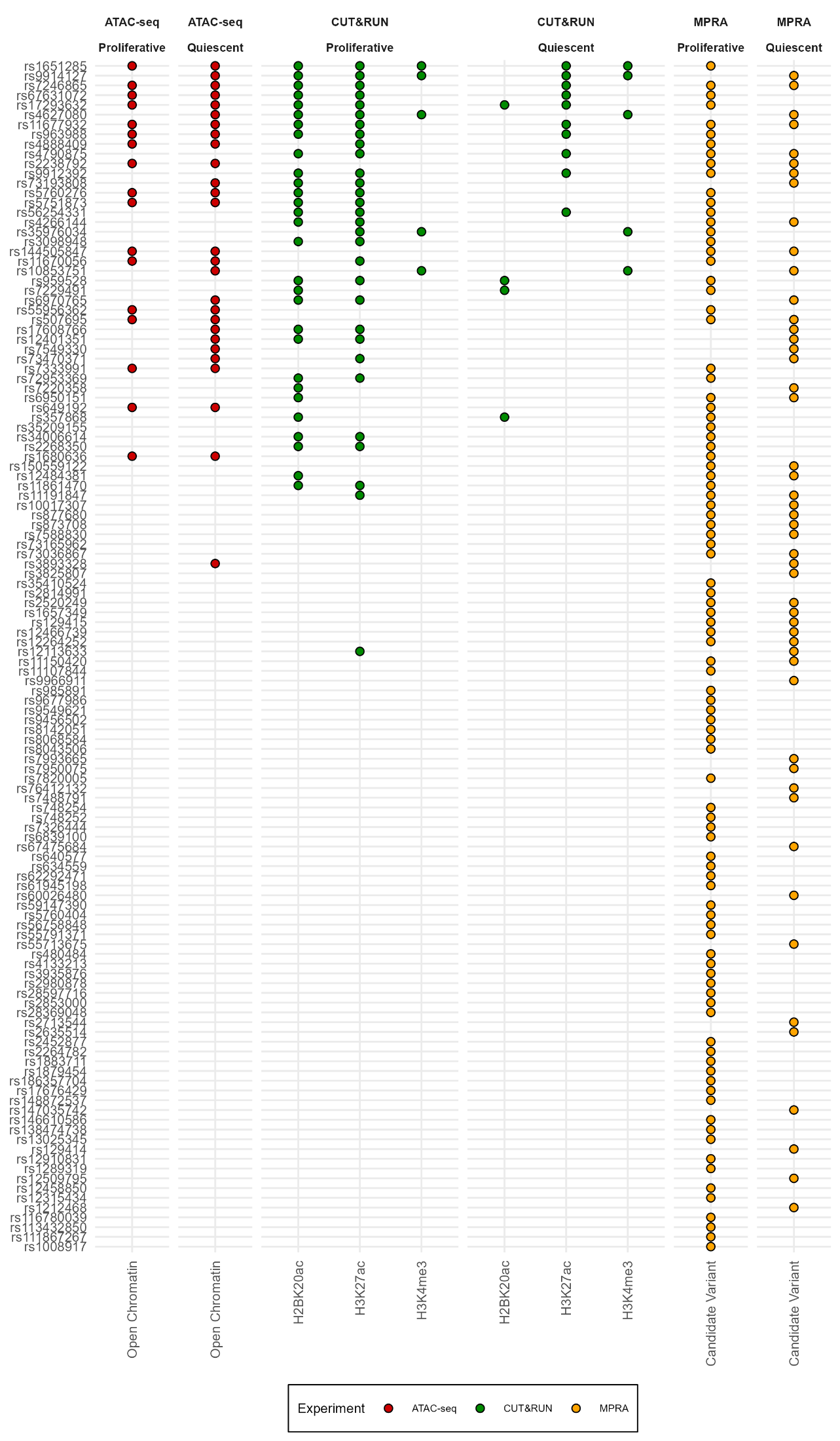

**Supplemental Figure 6. Functional annotation of lentiMPRA SNPs.** List of the lentiMPRA SNPs along with the epigenomics assays they appear in. Colored dots denote supporting evidence for the SNP in a given assay: red denotes that the SNP is present in a region of open chromatin (based on ATAC-seq), green denotes that the SNP is present in an enhancer or promoter region (based on CUT&RUN), and orange denotes that the SNP was identified as a SNP showing allelic imbalance in the lentiMPRA.

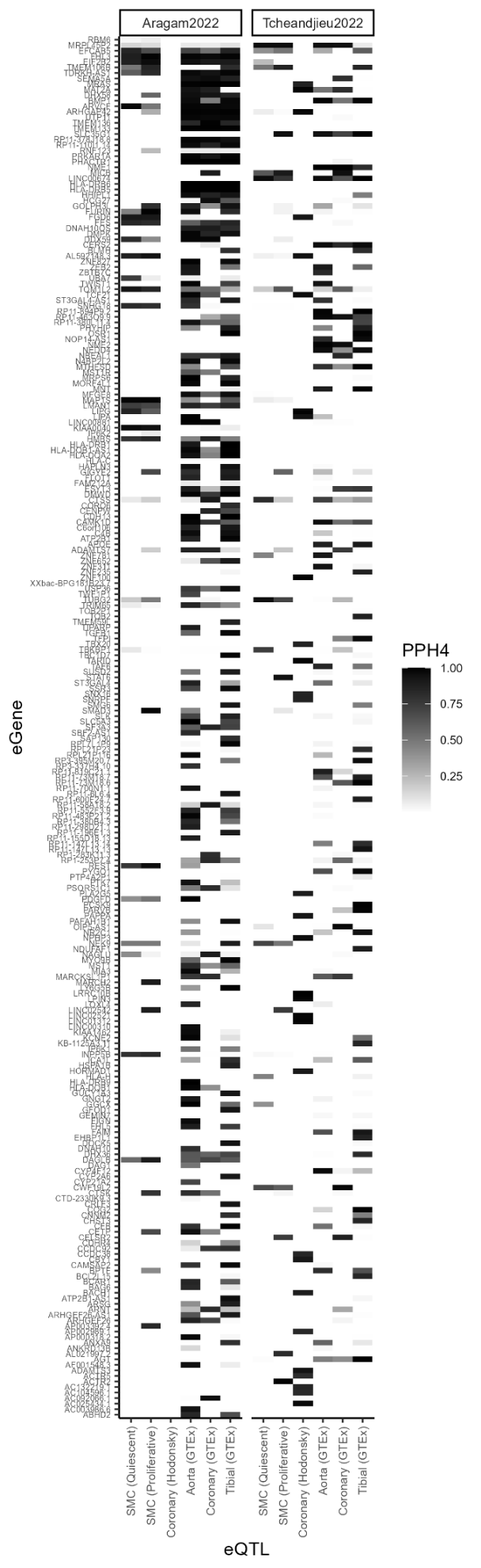

**Supplemental Figure 7. GWAS colocalization to eGene associations:** Each column represents a different eQTL study. Smooth muscle cell eQTL dataset is from reference 12; Coronary artery eQTL dataset is from reference 29; Aorta artery, tibial artery, and coronary artery eQTL data is from the GTEx project. The intensity of shading corresponds to posterior probability of a shared association at that locus. The darker color indicates a higher probability of colocalization. Each row represents one protein-coding gene with a PPH4 > 0.8 in at least one GWAS. Colocalization was performed in two different GWAS, shown in different panels.
